# Benchmarking metagenomic marine microbial growth prediction from codon usage bias and peak-to-trough ratios

**DOI:** 10.1101/786939

**Authors:** Andrew M. Long, Shengwei Hou, J. Cesar Ignacio-Espinoza, Jed A. Fuhrman

**Affiliations:** Department of Biological Sciences – Marine and Environmental Biology, University of Southern California, Los Angeles, CA, USA

**Keywords:** microbial growth rate, codon usage bias, CUB, peak-to-trough ratio, PTR

## Abstract

Growth rates are fundamental to all organisms and essential for characterizing microbial ecologies. Despite this, we do not know the instantaneous nor maximum growth rates of most naturally-occurring microorganisms. Recent reports indicate DNA replication rates can be estimated from metagenomic coverage, and maximum growth rates can be estimated from genomic characteristics. We tested these approaches with native marine bacteria (<0.6 um size fraction) as 10% inoculum grown in unamended virus-free seawater from the San Pedro Channel, California. This allowed microbial growth without grazing and with greatly reduced viral infection. At multiple time points up to 48 h, we sampled for cell abundances and metagenomic analyses. We generated 101 unique Metagenome-assembled genomes (MAGs) including α, β, and γ Proteobacteria, Flavobacteria, Actinobacteria, Verrucomicrobia, Marine Group A/SAR406, MGII archaea, and others. We tracked the growth of each as the fraction of total metagenomic reads mapped to each MAG normalized with length, completeness, and total cell counts. Some MAGs did not grow appreciably, but those we could estimate had growth rates ranging from 0.08 to 5.99 d^−1^; these are the first reported growth rates for several of the groups. These metagenome-determined growth rates, which often changed within experiments, were compared to (a) DNA replication estimates from the ‘peak-to-trough’ ratio (PTR) as determined by three different approaches, and (b) maximum growth rates predicted from codon usage bias (CUB). For the large majority of taxa, observed growth rates were not correlated to PTR indices (r ~ −0.26 - 0.08), with exceptions being rapidly growing *Oceanospirillales* and *Saccharospirillaceae* (r ~ 0.63 - 0.92). However, CUB was moderately well correlated to observed maximum growth rates (r = 0.57). This suggests that maximum growth rates can be reasonably well-estimated from genomic information alone, but current PTR approaches poorly predict actual growth of most marine planktonic bacteria in unamended seawater.

## Introduction

Growth rate is fundamental to an organism’s ecology and necessary to conceptually or mathematically model microbial systems. Therefore, many efforts have been made to estimate the growth rates of marine microbes. Historically, these growth rate estimates have used time-course incubations of pure cultures or mixed communities, often involving isotopic tracers of DNA or protein synthesis (reviewed in: [1]). These approaches provide valuable information on the growth of cultures in lab conditions, or regarding average growth in mixed communities. However, because they do not distinguish the contribution of individual phylogenetic groups to community-wide rates, they cannot be readily applied to native uncultivated microbes in a taxon by taxon manner in their natural habitats, which is often more germane to understanding their ecologies. To address individual rates, several culture-independent growth rate estimation methods have recently been developed that use intrinsic characteristics of microbial genomes and discrete metagenomic samples to estimate either its maximum growth rate or take a snapshot of each taxon’s growth rate.

These recently developed genome-based growth estimates take two fundamentally different approaches that generate fundamentally different answers: codon usage bias (CUB) and peak-to-trough ratio (PTR). The CUB approach aims to predict maximum growth rates while the PTR approaches seek to approximate instantaneous growth via estimating DNA replication rates. CUB is based on the physiological strategy of cells, in which there is a tendency of highly expressed genes to prefer one set of codons corresponding to the most abundant tRNAs in the cell, while all other genes are more likely to use alternative sets of codons (from the less abundant tRNAs) for the same amino acids. The degree of difference between the usage of abundant and alternative synonymous codons equates to the amount of bias present in any genome. This codon usage bias is stronger for cells that can grow faster and have a particularly high priority to make ribosomes to facilitate their fast growth. Viera-Silva and Rocha calculated the CUB of a wide range of prokaryotic taxa and found a strong correlation between observed maximum observed growth rates and CUB, which they leveraged to predict maximum growth rates by a multivariate approach [2]. Further, Kirchman found a strong correlation between observed and predicted maximum growth rates when applying this methodology to cultured marine taxa (including *Prochlorococcus*, SAR11, and others) in a recent review [1]. While CUB maximum growth estimators have been validated with pure cultures of organisms whose genomes are fully sequenced, it has yet to be tested with mixed natural communities, including uncultivated organisms growing on natural dissolved organic matter in seawater.

In contrast to CUB, PTR is an approach that is designed to work as an instantaneous measure of the actual growth rate of any prokaryote in any sample where genomic or metagenomic data are available. Thus, it can be remarkably powerful, and its attraction obvious. This method is based on the observation that prokaryotes generally have circular genomes that are replicated bidirectionally from a fixed origin to a fixed terminus (opposite the origin). When the microbial genomes are fragmented and sequenced, as is done in metagenomics, the PTR of the population represented by a genome could be calculated from recruiting metagenomic reads across that genome. Populations more rapidly growing and replicating their DNA will be expected to have more reads recruited to and near the origin of replication (‘peak’) compared to the terminus (‘trough’). Following a simple conceptual model, the slope of the resulting read recruitment sine curve should be reflective of the growth of the population, with steeper slopes indicating faster growth rates. The PTR method was first developed in practice by Korem *et al*. [3] for use with complete genomes and validated with *E. coli* growing in a chemostat. However, we do not have complete genomes for the large majority of prokaryotes in nature, and at best we often have partial genomes that are metagenomically assembled (MAGs). Such ‘genomes’ (actually genomic bins) are usually in many fragments and not only is the genomic order of those fragments unknown, but there is uncertainty about the locations of the origin and terminus. To address this problem, multiple methods modified the original PTR approach for use with incomplete genomes and MAGs. The first (iRep) was by Brown *et al*. [4], followed with a version for MAGs with low coverage (GRiD) by Emoila and Oh [5], and for MAGs with low coverage, low genomic completion, and high genomic redundancy (DEMIC) by Gao and Li [6]. While all three calculate PTR, one of their key differences is how they estimate the origin of replication and the terminus of a MAG. iRep uses coverage across overlapping genome fragments (windows) and then sorts the fragments from highest to lowest coverage to estimate the origin of replication and the terminus. Similarly, GRiD sorts genome fragments from highest to lowest but places fragments containing the *dnaA* gene at the origin and fragments with the *dif* gene near the terminus. Lastly, DEMIC infers the genome fragment placement using relative distances with a principal component analysis of fragment coverage in multiple samples. All of the PTR approaches have shown great promise when applied to pure cultures of various bacteria, using the data from either the Korem et al study or from other bacterial growth studies [4, 5, 6]. However, these approaches have not been validated in complex microbial communities such as those found in marine surface waters. Such communities potentially include highly non-uniform populations of individuals with different recent histories and with many co-occurring close relatives. Furthermore, many such organisms have slow growth rates, with division times much longer than the minimum time it takes to replicate a genome, and with individuals probably growing at a variable rate over time (as unpredictable resources and/or inhibitors change). The strategies by which such cells manage DNA synthesis and other cell components in preparation for cell division under natural dynamic conditions are not known and could make applying PTR to such cells challenging.

The purpose of this study is two-fold: 1) to estimate the growth rates of a broad variety of native planktonic marine microbes in natural dissolved organic matter *via* time-series incubations with unamended seawater and 2) to test and potentially validate the CUB and PTR approaches using metagenomic bins recovered from marine microbes growing in a complex community. In order to achieve this, we grew such bacteria in conditions meant to remove grazing and eliminate as much viral infection as possible, so that the observed growth would reflect the actual growth rates. From these, we used metagenomics to generate MAGs and assessed each MAG’s growth rate using the number of recruited reads normalized to direct cell counts, MAG completeness and cumulative contig length (equivalent to draft genome size) over the course of roughly 48 hours. The estimated growth rates were then compared using linear regression to CUB maximum growth rate predictions and PTR-derived growth rate indices.

## Methods and Materials

### Sampling and experimental design

Growth rate experiments were conducted in May and September of 2017. For each experiment, surface water was collected at 33°33’ N, 118°24’ W during the monthly sample collection of the San Pedro Ocean Time-series. Bulk samples (> 40 L) were collected on site and placed into coolers for transportation to USC. Upon arrival, bulk samples were first filtered through 80 μm nylon mesh (Sefar, Buffalo, NY, USA) and then sub-sampled into two pools. The first pool was pumped through Whatman^®^ 47 mm 0.6 μm track-etch PC filters (GE Life Sciences, Marlborough, MA, USA) twice with a goal of a grazer-free sample. The second pool was pumped through a Prep/Scale-TFF Cartridge 30 kD 2.5 ft^2^ (Millipore, Billerica, MA, USA) to remove viruses. These two pools were combined to dilute the 0.6 μm-filtered microbial communities to 9.8 % in May and 9.5 % in September. Duplicate 10 L samples were incubated in the dark in PC bottles at 17 □ in May and triplicate 10 L samples were incubated under the same conditions in September. Samples were taken at 0 h, 12 h, 24 h, and 48 h in May and 0 h, 11 h, 20 h, 37 h, and 44 h in September for DNA extraction and cell counts. Cells were counted with the SYBR green method described by Noble and Fuhrman [7] in May and a modified Acridine Orange Hobbie et al. [8] method in September, also described by Noble and Fuhrman [7].

### DNA extraction, sequencing, assembly and metagenome-assembled genome generation

For DNA extraction, water was pumped through 0.2 um Durapore Sterivex™ filter units. DNA was extracted from Sterivex™ filter units using an All-prep^®^ DNA/RNA minikit (Qiagen, Hilden, GR) with a modified protocol. Briefly, ~ 100 uL of combusted 0.1 mm glass beads (BioSpec Products, Bartlesville, OK, USA) were added directly to the Sterivex filter with lysis buffer from the All-prep^®^ kit and mixed on a vortex mixer (VWR model VM-3000) for 20 minutes at the maximum setting, the liquid was retrieved from the filters and the manufacturer’s protocol was followed thereafter. The resulting DNA was processed for sequencing using Ovation^®^ Ultra-low V2 DNA-Seq Library Preparation kits (NuGen, Tecan Genomics, Redwood City, CA, USA) with the manufacturer’s protocol using 100 ng of starting DNA and 9 PCR cycles. DNA was sequenced on an Illumina HiSeq platform at the USC UPC Core Sequencing Facility (Los Angeles, CA, USA) using 2×250 chemistries.

The computer programs atropos v1.1.18 [9] and sickle v1.33 [10] were used to remove adaptor sequences and bases with quality scores below 25, which was then verified with fastqc v0.11.5 [11]. All samples were assembled individually with metaSPAdes v3.12.0 [12] with a custom kmer set (-k 21,33,55,77,99,127) under the following subsampling regime: first 1% of the reads, 1.5%, 2%, 5%, 10%, 20% were assembled separately, and then 5%, 10%, 20%, 33%, and 50% were assembled sequentially without replacement of assembled reads (i.e., 5% of the reads were assembled and then 10% of the remaining reads were assembled without including reads that were assembled from the 5% step and so forth). These subassemblies were sorted into two sets based on contig length cutoff (2 kb) using seqkit v0.3.4.1 [13]. Those contigs shorter than 2 kb were assembled using Newbler v2.9 [14] with a minimum identity cutoff 0.98 (-mi 98) and a minimum overlap 80 nt (-ml 80), the resulting assembled contigs (>= 2kb) were then combined with the longer contig set. All these contigs longer than 2 kb were further coassembled using minimus2 from the AMOS v3.1.0 toolkit [15] with a minimum identity 0.98 (-D MINID=98) and a minimum length cutoff 200 nt (-D OVERLAP=200). The co-assembled contigs were de-replicated using cd-hit v4.6.8 [16] with a 0.98 identity cutoff (-c 0.98), the dereplicated contigs were renamed and were used as references for read recruitment with bwa v0.7.15 [17] and the following metagenomic binning. MetaWRAP v1.1 [18] was used to bin contigs via MetaBAT v2.12.1 [19], CONCOCT v1.0.0 [20], and MaxBin2 [21] with a minimum length cutoff of 2 kb. The resulting bins were further refined within MetaWRAP without filtering for completion (-c 0) and allowing high contamination (-x 10000). Anvi’o v5.1.0 [22] was also applied to bin contigs > 5kb using CONCOCT proceeded by manual refinement with redundancy cut-offs of 2.5% for MAGs with 50 - 75 % completeness, 5 % for MAGs with 75 - 90 % completeness and 10 % for MAGs with > 90 % completeness. In addition, Vamb v1.0.1 [23] and BinSanity v0.2.8 [24] were used to bin contigs with a minimum length cutoff of 4 kb. All bins generated by the 6 binners and the MetaWRAP refined bins were further refined using DAS_Tool v1.1.1 [25] with custom penalty parameters (--duplicate_penalty 0.4, -- megabin_penalty 0.4) and a score threshold of 0.3. All the DAS_Tool refined bins were further refined manually using anvi’o v5.1.0 [22]. MAGs with at least 50% completion and fewer than 5% redundancy or at least 90% complete and fewer than 8% redundancy according to anvi’o were retained for further analysis. The GTDB taxonomic information of manually refined bins were predicted using GTDBTk v0.1.3 [26] and compared to NCBI taxonomy commonly used to provide more context to the previous literature where appropriate.

### Phylogenomic analysis

The bacterial and archaeal phylogenomic trees were constructed using GToTree v1.1.3 [27] and RAxML-NG v0.8.1 [28]. Briefly, GToTree uses Prodigal v2.6.3 [29] to predict the coding regions of the MAGs and uses HMMER v3.2.1 [30] to search for 74 bacterial and 76 archaeal universal single-copy marker genes. Then, these marker genes are concatenated and aligned using MUSCLE v3.8 [31]. Next, the alignments were trimmed using Trimal v1.4 [32] with the heuristic “-automated1” method. Both the bacterial and archaeal phylogenomic trees were constructed using RAxML-NG based on the GToTree produced trimmed alignments. RAxML-NG was run with 10 randomized parsimony starting trees (--tree pars{10}), a fixed empirical substitution matrix, a discrete Gamma model with 8 categories of rate heterogeneity, empirical amino acid state frequencies estimated from the sequence alignment (--model LG+G8+F), and was performed for 200 non-parametric bootstrap replicates (--bs-trees 200).

### Growth rate estimations

First, the relative abundance for each MAG was calculated from the number of reads that mapped to that specific MAG using bwa v0.7.15 [17] under the default parameters, which was then corrected with the completeness information from anvi’o v5.1.0 [22] and the length (in bp) of the MAG. This theoretical number of reads that map to each MAG was then divided by the total number of reads in each sample to calculate the relative abundance. The completeness and genome length adjusted relative abundance of each MAG was multiplied by the cell count at each time point to estimate the absolute abundance for each MAG.

The differences between the natural logarithm-scaled cell abundances of two time points were divided by the time interval (in hours) and multiplied by 24 to estimate the growth rate per day. The observed maximum growth rates were taken from the highest estimates between the following time scales: 0 h - 36 h and 18 h - 42 h for May; 0 h - 20 h, 11 h - 37 h, and 20 h - 44 h for September.

### Growth rate indices: codon usage bias and peak-to-trough ratio

The maximum growth rate of each MAG was predicted using a customized growthpred v1.0.8 (available at https://hub.docker.com/r/shengwei/growthpred) in metagenome mode (-m) and with universal codons (-c 0). Blast-retrieved ribosomal protein sequences were used as the highly expressed genes (-b) and compared to all the coding sequences of each MAG (-g).

PTR indices were calculated using iRep v1.10 [4], GRiD v1.3 [5], and DEMIC v1.0.2 [6]. iRep and GRiD were calculated for all MAGs > 75% complete and DEMIC was calculated for every MAG. Briefly, the mapping information for each MAG was extracted from the previously generated bam files, then the iRep and GRiD indices were calculated based on the aligned paired reads, determined for: (1) the entire metagenomic dataset for that sample, and (2) reads from metagenomic fragments with specific ranges of insert sizes (100-350, 200-450, 300-550, 400-650) to evaluate the effect of different insert sizes on the performance of PTR indices. In addition, GRiD was also run under the default parameters using each individual MAG as input. DEMIC was run under the default parameters using sam files generated with bowtie2 v2.3.0 [33].

### Statistical comparisons

To compare observed growth rates with predicted maximum growth rates from growthpred, we used the highest overall growth rate from each taxon based on three time points. For the PTR indices, the growth rates estimated from the time points adjacent to the point at which the PTR index was calculated were used for comparisons. For instance, if the PTR index was calculated from time point 1, the growth rate it was compared to was estimated using time points 0, 1, and 2. For all comparisons, linear regressions were calculated in R version 3.5.3 using the package ggplot2 and Pearson correlation coefficients and p-values were calculated using the package Hmisc [34]. When a MAG had no apparent growth during the time frame, their corresponding growthpred or PTR index values were removed before statistical analyses. Further, outliers as calculated using Tukey’s fences in PTR index values and their corresponding growth rates were excluded from statistical analyses.

### Data availability

Raw reads and sample information have been submitted to NCBI under project ID PRJNA551656, the dereplicated assemblies and manually curated MAGs have been deposited at https://doi.org/10.6084/m9.figshare.9730628.

## Results

### Metagenome-assembled genomes

We recovered 101 MAGs that passed our quality criteria. The MAGs are on average 70 % complete, 3 % redundant, and cover most major groups of marine planktonic bacteria as well as MGII Euryarchaeota (Fig 2). Several MAGs were from groups that have no cultivated representatives (Supplementary Table 1).

**Figure 1.**
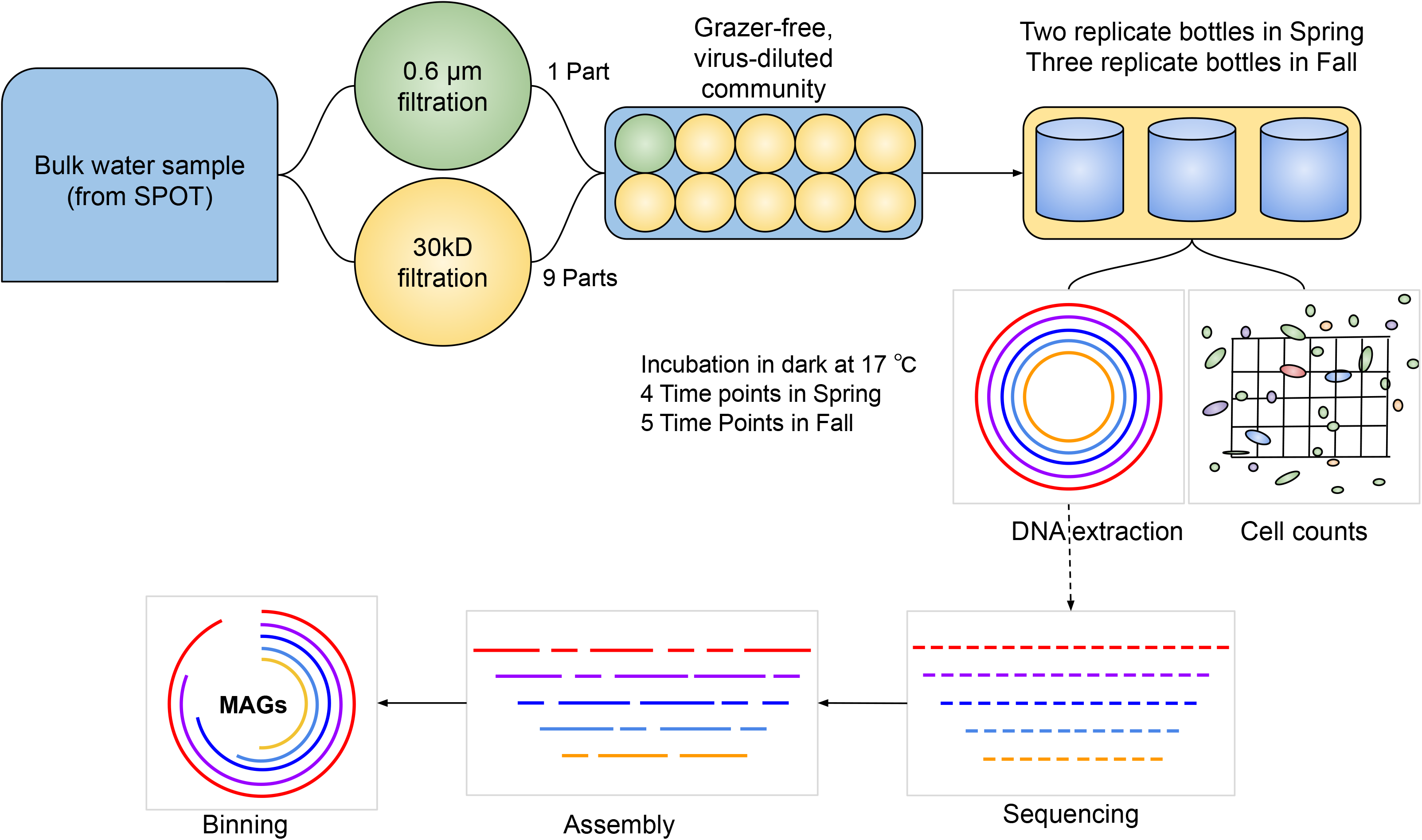
Flowchart of experimental design and data analysis

**Figure 2.**
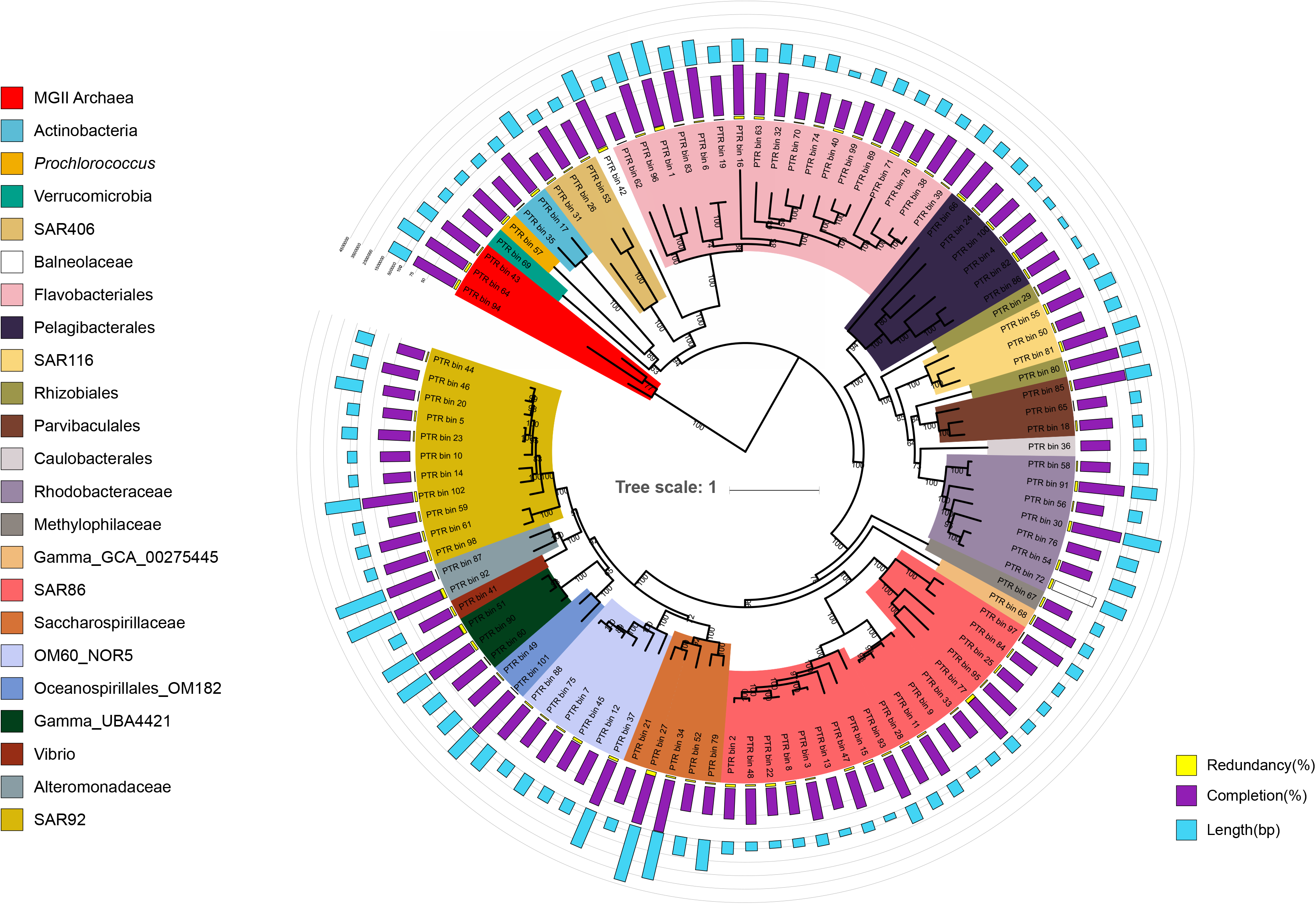
Phlyogenomic tree of metagenomic-assembled genomes, with information on G+C content, predicted genome size, completion and redundancy.

### Growth rate estimation from Metagenome-assembled genomes

The overall range of highest estimated growth rates for all MAGs with detectable growth was 0.08 – 5.99 d^−1^ (Fig 3). *Oceanospirillales Saccharospirillaceae* MAGs had the highest observed growth rates, of 3.17 – 5.99 d^−1^. Among *Pelgibacterales*, the highest observed growth rates ranged from 0.40 – 0.58 d^−1^. The majority (60 of 101) of MAGs had higher growth rates in the September incubation experiment than in May (Fig 3). However, many of the fastest growing taxa grew faster in the May incubation, such as MAGs affiliated to SAR92 and Flavobacteriales.

**Figure 3.**
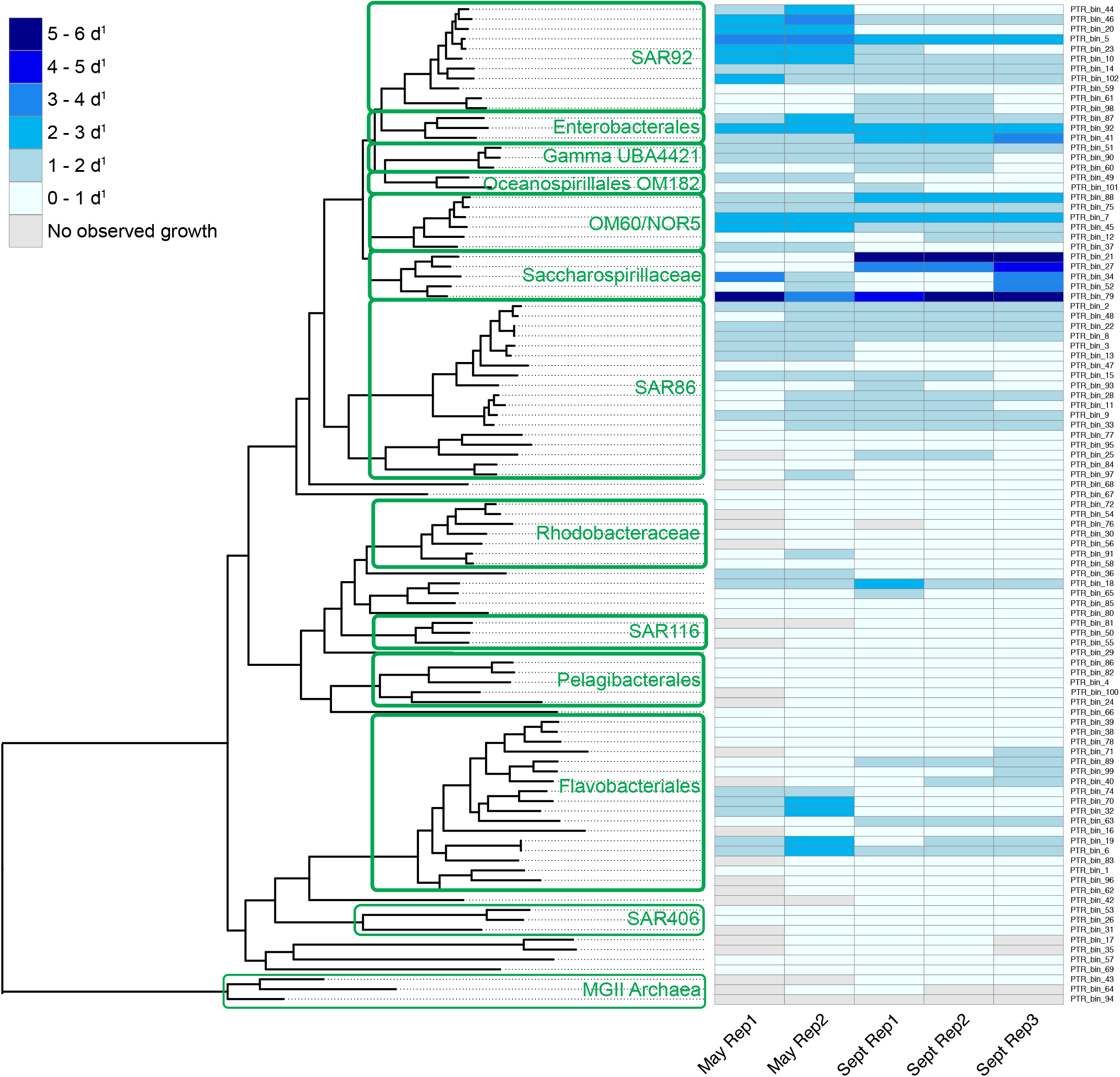
Phylogenomic tree with associated heatmap showing highest observed growth for each taxon in each experiment and replicate.

### Highest observed growth rates compared to codon usage bias max growth rates

Growthpred predicted a total range of maximum growth rates based on CUB of 0.40 – 16.47 d^−1^ (Supplementary Table 1). The highest predicted maximum growth rates were from *Oceanospirillales Saccharospirillaceae, Vibrionaceae*, and *Alteromonadaceae* MAGs and the lowest were from *Betaproteobacteria, Pelagibacterales*, and SAR406 MAGs. Seventy-four of the 101 MAGs had a lower observed maximum growth rate during the experiments than a predicted maximum growth rate (Fig 4). Pearson correlation analysis with predicted max growth rates and highest observed growth rates of all MAGs found a good correlation (r = 0.57, p < 0.00001, n = 101), especially considering the observed growth is unlikely to be at the maximum actual rate.

**Figure 4.**
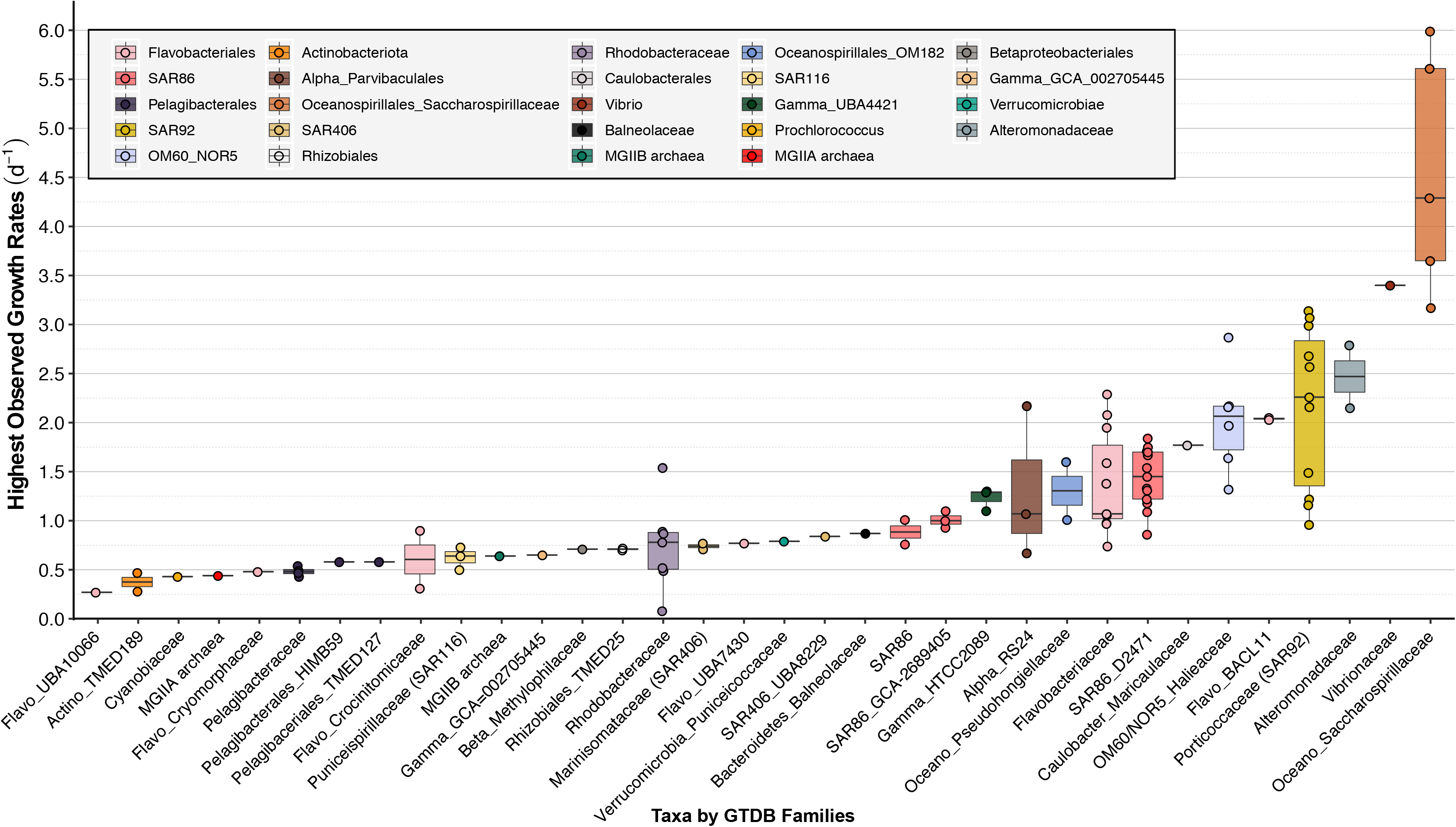
Box plot of highest observed growth rate for each MAG, grouped by taxonomic family. The boxes are drawn from the 25^th^ to 75^th^ quantile and center line of each box is the median. Whiskers indicate the smallest and largest value for each taxonomic family.

**Figure 5.**
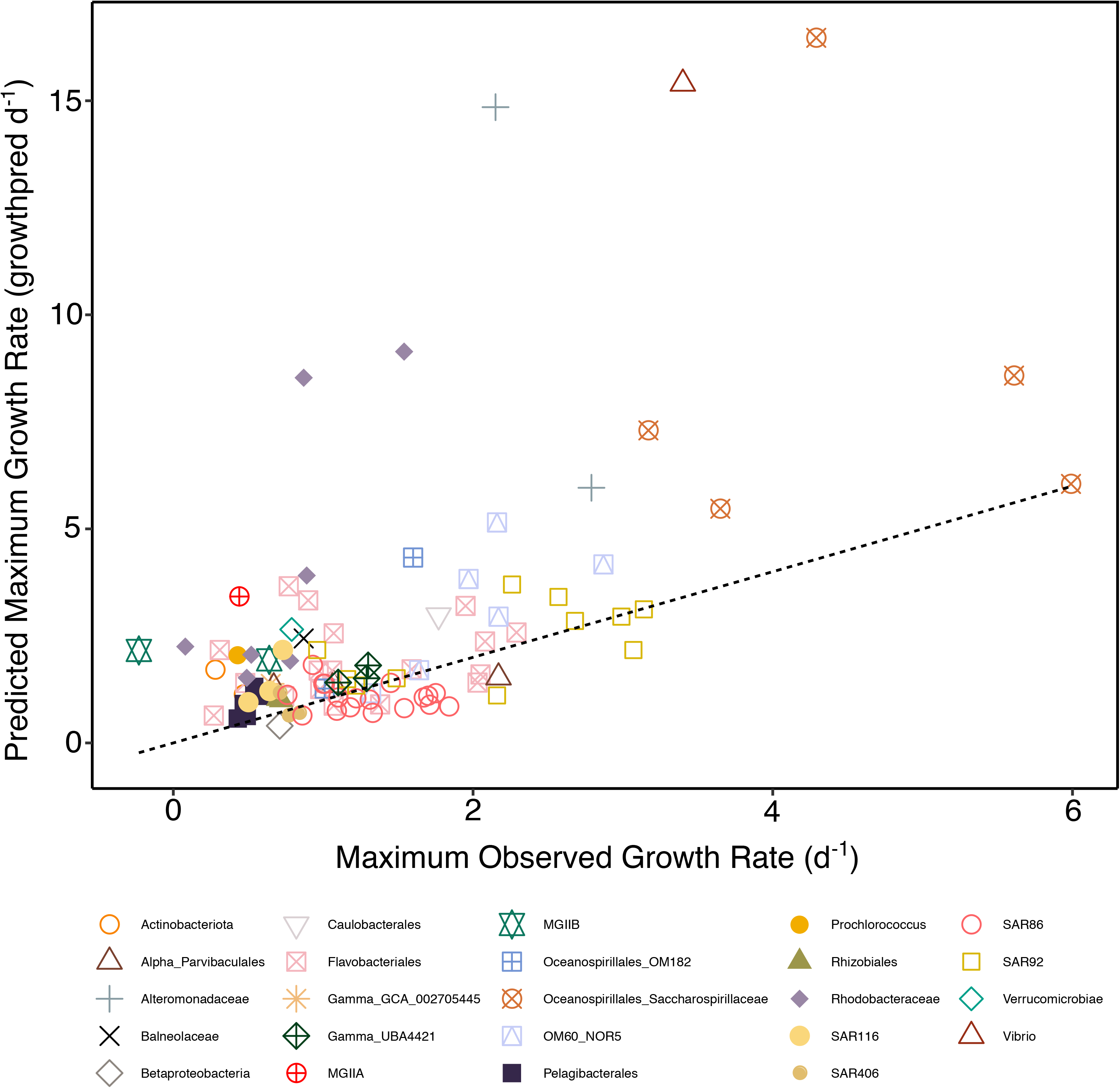
Highest observed growth rate over three time points plotted against predicted maximum growth rates from codon-usage bias predictor (growthpred). Dotted line is x = y to illustrate which MAGs have a higher predicted maximum growth rate than their highest observed growth rate.

**Figure 6.**
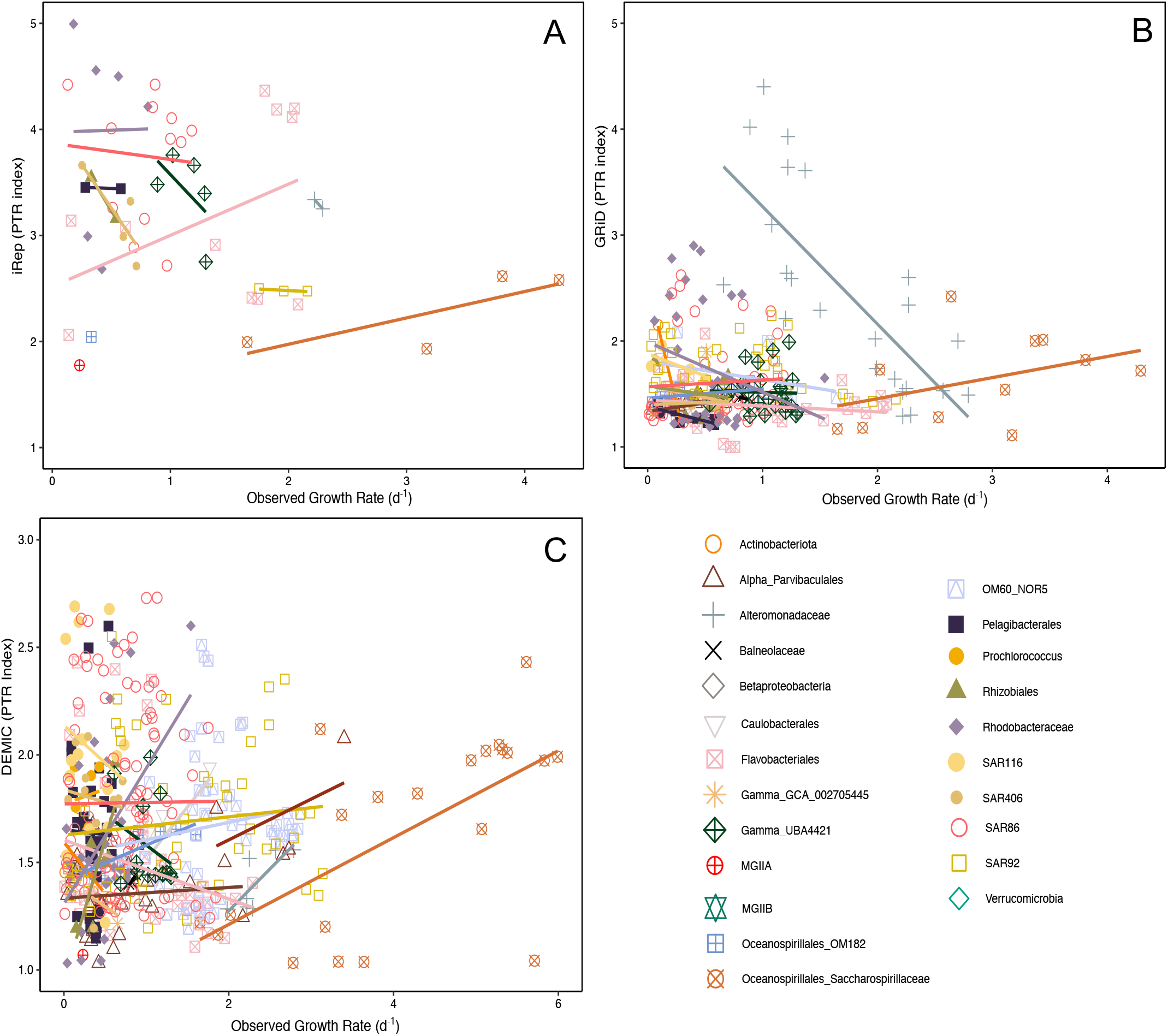
Observed growth rate over three points against peak-to-trough ratio indices: A) iRep B) GRiD C) DEMIC. Lines are linear regressions for each taxon with more than two observations of both growth rates and PTR index.

### Observed growth rates compared to peak-to-trough ratio indices

The range of the three PTR indices were 1.69 - 4.99 (iRep, mean = 3.15), 1 - 4.4 (GRiD, mean = 1.63), and 1.01 - 12.82 (DEMIC, mean = 1.99). The correlations between PTR indices and observed growth rates of all taxa combined were either negative (iRep, r = −0.27, p-value = 0.053, n = 52) or weak (GRiD, r = 0.077, p-value = 0.20, n = 273; DEMIC, r = 0.072, p-value = 0.13, n = 446). When comparing the observed growth rates and PTR values taxon-by-taxon, the majority of taxa had either negative or weak correlations (Supplementary Table 2). However, for every PTR index, *Oceanospirillales Saccharospirillaceae* had a good correlation between observed growth rates and PTR index (iRep, r = 0.78, p-value = 0.22, n = 4; GRiD, r = 0.40, p-value = 0.22, n = 11; DEMIC, r = 0.63, p-value = 0.0022, n = 21). *Oceanospirillales* OM182 also had good correlations between observed growth rates and GRiD (r = 0.49, p-value = 0.18, n = 9), and DEMIC (r = 0.92, p-value = 0.004, n = 9). Only *Oceanospirillales Saccharospirillaceae* and *Oceanospirillales* OM182 MAGs had good, positive, and significant correlations with an alpha of 0.01 between observed growth rates and any PTR index (DEMIC in this case).

## Discussion

### Bacterial growth rates of uncultivated clades

The MAG-derived growth rate estimation approach allowed us to obtain growth information from clades without cultured representatives such as SAR406, SAR86, and MGII Euryarchaeota. The growth rates estimated for SAR406 (0.29 – 0.39 d^−1^) were similar to those of several other heterotrophs, such as MAGs belonging to *Pelagibacterales* and SAR116. As expected, the observed growth rates were lower but close in value to the CUB predicted maximum growth rates. This result is interesting because SAR406 has usually been found at higher abundances in suboxic and hypoxic environments, leading researchers to think they preferred such conditions [35], and our conditions were fully aerobic. The range of predicted maximum growth rates (0.64 - 1.17 d^−1^) suggests an oligotrophic lifestyle. Additionally, SAR86 MAGs also had a low range of both observed (0.53 - 1.84 d^−1^) and predicted (0.64 - 1.82 d^−1^) maximum growth rates. Most previous studies examining SAR86 growth also suggest low activity (e.g., [36–38]), except for a study in the coastal North Sea [39]. However, as SAR86 is thought to have at least three distinct subclades [40–41], these previously published results may not be directly applicable to the SAR86 MAGs recovered in our study.

Two groups of MGII Euryarchaeota were present in our experiments: MGIIa and MGIIb. MGIIa are typically found in higher abundances in surface waters [42], whereas MGIIb are more prevalent in deeper waters (e.g., [43–45]), though found in our surface sample. Despite these relatively low observed growth rates (0.19 – 0.44 d^−1^), all three MGII MAGs had predicted maximum growth rates based on CUB near or over 2 d^−1^. This suggests that MGII may be quite active in certain environments, as we previously reported for SPOT where MGII 16S rRNA sequences comprised over 40% of the microbial community (higher than SAR11) on one post-spring-bloom day [47]. Further, MGII have also been found to be abundant in Monterey Bay, CA [48] and the Northern Gulf of Mexico “Dead Zone”, where they were found to be over 10% of the total microbial community [35].

In addition to taxa without cultivated representatives, we were able to obtain growth information on a number of representatives of previously cultured groups that have important ecological roles, such as *Pelagibacterales*. The ranges of both observed and predicted growth rates for *Pelagibacterales* MAGs were within the range of previously published observed growth rates (0.4 - 0.6 d^−1^; reviewed in [1]).

### Codon usage bias maximum growth rate predictions

Growthpred, which predicts the maximum growth rate from CUB [2], had a significant statistical relationship with the MAG-derived observed maximum growth rate estimates. While Kirchman [1] applied this methodology to cultured organisms with complete genomes and found a similarly strong relationship, this is the first validation of the method using MAGs and growth rates obtained in a mixed microbial community. The good relationship between the highest observed growth rates and predicted maximum growth rates suggests that growthpred works reasonably well for prokaryotes that can be binned into a high-quality MAG (here defined as > 50 % completeness and < 5 % redundancy according to single-copy genes), at least within the range of growth rates we observed. This includes MAGs belonging to clades without cultured organisms, such as those detailed in the above section (SAR406, SAR86, and MGII Euryarchaeota). In addition, organisms previously known to be adapted to oligotrophic environments, such as *Pelagibacterales*, had low predicted maximum growth rates (0.57 - 1.31 d^−1^) and organisms with high growth rates in previous studies, such as Vibrio (up to 14 d^−1^ in [49]) had high predicted maximum growth rates (15.4 d^−1^). These results confirm Kirchman’s [1] conclusion from cultured marine bacteria that growthpred can be applied reasonably well for the estimation of maximum growth rates for many fast and slow-growing marine bacteria, and we extend these results to at least one archaeal clade, and importantly to organisms that have not yet been cultured. However, some slow-growing organisms, like *Prochlorococcus*, had a much higher predicted maximum growth (2.05 d^−1^) than any previous estimation of their growth rate (0.1 - 1 d^−1^; e.g., [50–52]). Thus, while the relationship between observed growth rates and predicted growth rates across all taxa is strong, some taxa may have their growth potential overestimated with CUB. Others, as evidenced by the 27 of 101 MAGs with higher observed growth rates than predicted maximum growth rates, apparently have their growth potential underestimated. It is possible that growthpred might be modified to more accurately predict maximum rates, perhaps by considering data from this and other similar experiments.

### Peak-to-trough ratio in situ growth indices

The lack of a strong positive relationship between observed growth rates and all the variants of the PTR growth indices across all taxa suggests that these methods need rigorous improvement in order to use with MAGs in complex natural microbial communities. However, *Oceanospirillales* MAGs, which were some of the fastest growing MAGs in the experiments, had strong positive correlations between growth rates and PTR indices. *Oceanospirillales* MAGs started off at very low abundances relative to the rest of the community and grew swiftly to become one of the dominant taxa at the end of the experiment. Thus, PTR in natural mixed populations may be more suitable for rapid than for slow growth. One consideration is that this fast growth from a small initial population may have made *Oceanospirillales* MAGs more genomically clonal than the organisms that started off at high abundance and grew slowly. Considering that one of the challenges in applying PTR in mixed natural communities is cross-recruitment from close relatives when mapping reads, then it is reasonable that a more clonal population would reduce the noise generated from such cross-recruitment. Such co-occurrence of many close relatives is a characteristic frequently observed in natural populations (e.g., [53–54]). This also raises the question about the extent to which MAGs represent an amalgamation of closely related strains with potentially different growth rates. This may then lead to irregular and noisy read recruitment and thus PTR index calculation. For example, small mutations that alter growth might not be represented even in the most rigorously generated MAG, or MAG generation steps may merge related strains that might legitimately be considered part of a “population” but grow differentially. PTR indices from such strains would probably represent an average between merged strains but would be noisier than if done strain-by-strain. To examine this issue, we compared an alternative to our regular MAG generation protocol that eliminated overlap assembly and thus was likely to merge fewer close strains, and those results yielded no better relationships between growth and PTR indices (Supplemental Figure 1).

Besides the issue of clonality, there are other challenges for the application of PTR. Another involves within-population physiological uniformity – notably, PTR implicitly assumes individuals of a population are growing at the same rate. But there is no reason to necessarily expect such uniformity in natural populations where even genetically uniform individuals could have had very different histories and conditions at the time of sampling, i.e. some may have been in more rich microenvironments than others, or be virus-infected, and hence may be at different physiological states and growth rates. Other potential problems with PTR interpretation may relate to our lack of knowledge on how diverse slowly growing natural populations replicate their genomes in nutrient-limited, spatially patchy, and time-variable environments. It would already be a challenge to understand PTR-growth relationships under steady very slow growth conditions, i.e. does replication occur slowly, proportionately to growth, or does it start and stop many times, or do cells save up precursors and replicate the whole genome in only a small fraction of a generation? We expect this becomes particularly complicated when cells experience very different conditions during one generation, as might be expected in patchy natural environments. There is also the question of synchronization, common in phototrophs (diel cycles) and reported for at least some physiological processes even in oligotrophic heterotrophs like SAR11 [55].

Metagenomes today are most commonly generated with library preparation kits that involve a PCR-amplification step (i.e. linker amplified shotgun libraries), including this study and the dataset used to test all PTR indices [3]. It has been recently recognized that this amplification step alters the relative abundance of metagenomic reads compared to the original DNA, specifically yielding a small-insert bias due to amplifying and sequencing small inserts more readily than larger inserts [56]. This bias produces artifacts in quantitative read-mapping, potentially altering PTR index values, or at least introducing considerable noise in the PTR estimation. We considered this possibility and tried to avoid this artifact by sub-setting the read-mapping files according to insert sizes (100 - 350, 200 - 450, 300 - 550, 400 - 650). We found that when we did so, PTR index values trended lower for the same MAG when using shorter insert sizes (Supplemental Table 3). However, the lowered PTR index values still produced weak and negative correlations when compared to observed growth rates and did not improve the PTR-based predictions of growth (Supplemental Table 2).

In conclusion, metagenomic data can provide useful growth information for organisms represented by MAGs. CUB maximum growth rate predictors generally worked well, but PTR-based approaches usually did not, except for some of the fastest growing taxa. Looking forward, the CUB method could possibly be improved by an updated training dataset that considers more slowly growing organisms, many of which are now in culture. Other alterations to the CUB approach may include consideration of other genes that are known to be highly expressed in addition to ribosomal proteins. It is also possible that improvements in metagenomic assembly, binning, and read mapping, as well as better information on DNA replication strategies in diverse nutrient-limited slow-growing populations, may improve prospects for PTR-based growth estimates and other related approaches.

## Acknowledgements

We are very grateful to the Captain and crew of the Yellowfin and to all of the members of the Fuhrman lab for their input when discussing this research. This work was supported by the Simons Collaboration on Computational Biogeochemical Modeling of Marine Ecosystems/CBIOMES grant 549943 to JF, the Gordon and Betty Moore Foundation Marine Microbiology Initiative grant 3779 and NSF grant OCE1737409

---

Supplemental Figure 1. Observed growth rate over three points from MAGs generated without overlap-assemblers against peak-to-trough ratio indices. Lines are linear regressions for each taxon with more than two observations of both growth rates and PTR index.

