## Supplementary figures and images for "Benchmarking metagenomic marine microbial growth prediction from codon usage bias and peak-to-trough ratios"

### Supplemental Figure 1

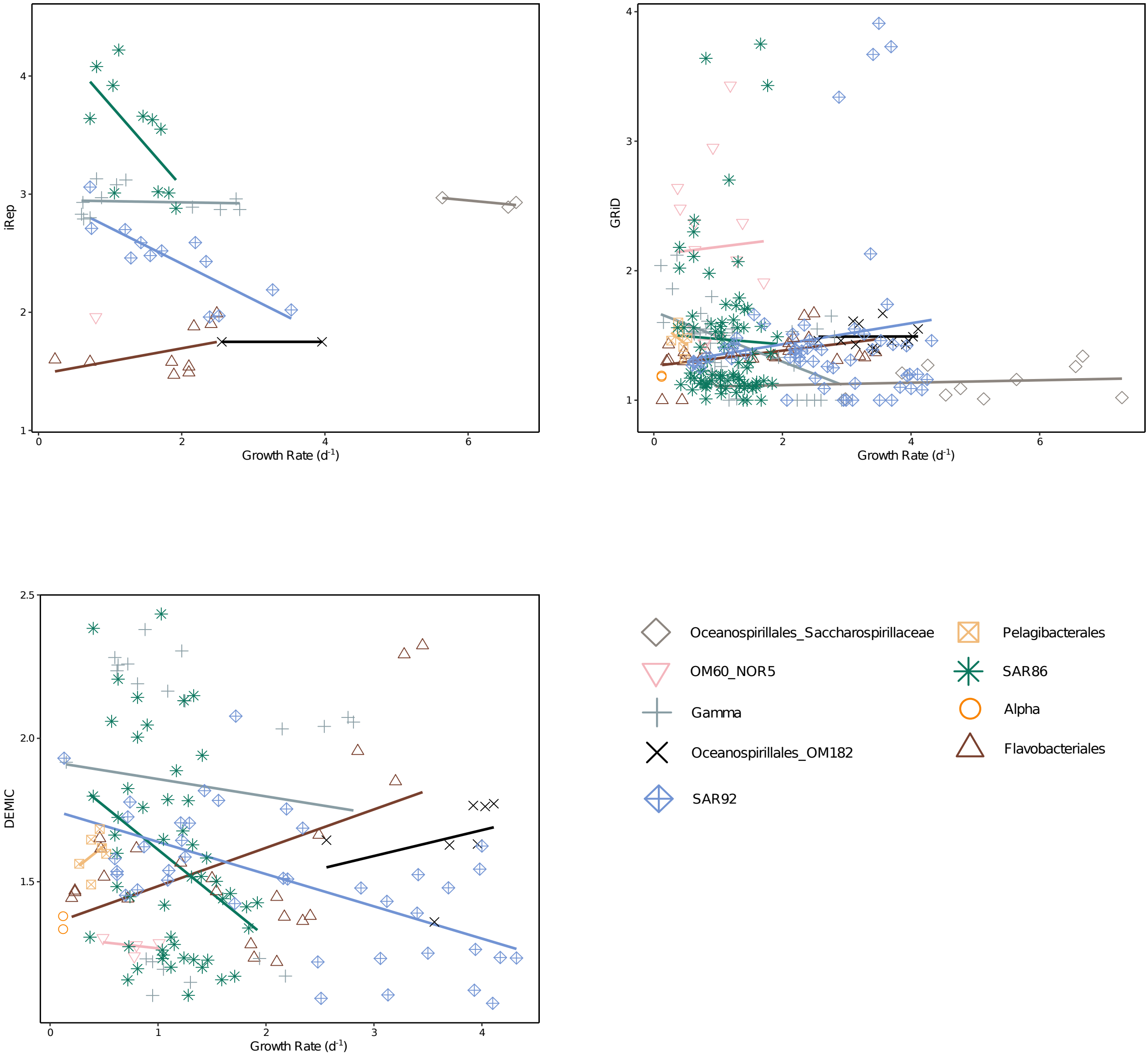
